# Reprogramming of larval locomotory pattern during baculovirus-induced host manipulation

**DOI:** 10.1101/2020.05.13.092627

**Authors:** Hiroyuki Hikida, Susumu Katsuma

## Abstract

Many parasites manipulate host behaviour to enhance their transmission. Baculovirus induces enhanced locomotory activity (ELA) combined with subsequent climbing behaviour in lepidopteran larvae, which facilitates viral dispersal. Previously, larval locomotion during ELA was summarized as the distance travelled for a few minutes at several time points. However, as ELA continues for several hours, these methods are unlikely to evaluate larval locomotion precisely during ELA. We developed a novel method to continuously trace the locomotion of *Bombyx mori* larvae using time-lapse imaging. This method successfully quantified the locomotory activities of larvae infected with *Bombyx mori nucleopolyhedrovirus* (BmNPV) for 24 h. We found that both mock- and BmNPV-infected larvae periodically repeated moving and pausing with a similar frequency. In contrast, BmNPV-infected larvae showed fast and long-lasting locomotion compared with mock-infected larvae, which resulted in longer locomotory distances in infected larvae. Moreover, BmNPV-infected larvae exhibited biphasic behaviour. Initially, BmNPV-infected larvae showed longer locomotory distances, but the locomotory pattern was similar to mock-infected larvae. However, during the second phase, the locomotory pattern was drastically altered, with an extremely larger locomotory area. These results indicate that BmNPV reprograms host locomotory pattern, which is a turning point for the process of BmNPV-induced host manipulation.

## 1. Introduction

Manipulation of host behaviour is a strategy commonly used by parasites to enhance their proliferation in the environment [1]. For example, hairworms cause crickets to jump into the water, where the hairworms can search for their mates [2], and *Ophiocordyceps* fungi make ants to bite onto vegetation at an elevated position, which is thought to enable the fungi to disperse their spores [3]. Baculovirus is an insect-specific large DNA virus, which mainly infects lepidopteran larvae and induces dramatic alternation to host behaviour, which is known as *Wipfelkrankheit* (tree-top disease in German) [4]. Baculovirus-infected larvae exhibit horizontal hyperactivity, which is called enhanced locomotory activity (ELA), followed by vertical movement, which is known as climbing behaviour (CB). These virus-induced host manipulations finally lead the larvae to the upper tree foliage, where they die. Baculovirus encodes cathepsin and chitinase in its genome, which cooperate to liquefy the larval body at a late stage of infection [5,6]. The combination of larval death at an elevated position and liquefaction of their body is thought to facilitate viral dispersal with rainfall or avian feeding [4,7].

Baculovirus-induced host manipulation was first quantified by Goulson in 1998 using Mamestra brassicae multiple nucleopolyhedrovirus (MbMNPV) and *Mamestra brassicae* larvae. The study demonstrated that MbMNPV-infected larvae show higher horizontal dispersal and die at an elevated position [4]. Recent studies identified baculovirus-encoding genes, *protein tyrosine phosphatase* (*ptp*) and *ecdysteroid UDP-glucosyl transferase* (*egt*), as factors for inducing larval locomotion during ELA and CB, respectively [8,9]. The deletion of *ptp* in Bombyx mori nucleopolyhedrovirus (BmNPV) and Autographa californica multiple nucleopolyhedrovirus (AcMNPV) impairs the horizontal locomotory activity of infected *Bombyx mori* and *Spodoptera exigua* larvae, respectively [8,10,11]. As *ptp* deletion affects ELA but not climbing height during AcMNPV infection, ELA and CB are thought to be governed by different mechanisms [12]. The deletion of *egt* in Lymantria dispar multiple nucleopolyhedrovirus and Spodoptera exigua multiple nucleopolyhedrovirus mitigates the climbing height of *Lymantria dispar* and *S. exigua* larvae, respectively, although the enhancement of larval activity by *egt* was not observed in BmNPV- and AcMNPV-infected larvae [9,13–15]. A recent study further showed that miRNA-mediated hormonal regulation affects the climbing height of *Helicoverpa armigera* larvae infected with Helicoverpa armigera single nucleopolyhedrovirus, indicating that hormonal cues trigger CB [16]. However, these studies assessed the larval locomotory level using a few sampling points of usually once per several hours. As ELA and CB are continuous events that occur for hours or days, these methods only evaluated a part of the baculovirus-induced host manipulation.

We developed a novel method to continuously observe the locomotion of BmNPV-infected *B. mori* larvae using short-interval time-lapse recording for 24 h, followed by a video analysis to trace larval locomotion. *B. mori* has been highly domesticated for silk production, so the species is generally inactive [17–19]. In contrast, they exhibit vigorous locomotion during BmNPV infection, which makes the species an optimal model to measure the change in locomotory activity following baculovirus infection [8]. We quantified the horizontal locomotion of BmNPV- and mock-infected larvae in cell culture dishes. Our method successfully traced larval locomotion at a 3 s resolution. It revealed that BmNPV-infected larvae periodically showed moving and pausing activity at a similar frequency to mock-infected larvae, but their locomotion was faster and longer-lasting than that of mock-infected larvae. We also found that the locomotion of both mock- and BmNPV-infected larvae was restricted around food at the early phase of observation. In contrast, the locomotory pattern of BmNPV-infected larvae was altered drastically at the late phase of observation, and their locomotory area became extremely larger. These results provide novel insight into the mechanisms of BmNPV-induced host manipulation system.

## 2. Materials and Methods

### (a) Insects, cell lines, and viruses

*B. mori* larvae (Kinshu × Showa) were reared as previously described [20]. BmN-4 cells were cultured in TC-100 medium supplemented with 10% fetal bovine serum at 26◻. The T3 strain was used as the wild-type BmNPV [21] and propagated in BmN-4 cells. Viral titre was determined by a plaque assay method using BmN-4 cells.

### (b) Locomotion assays

Fourth-instar larvae were starved for several hours and then injected intrahaemocoelically with a viral suspension containing 1 × 10^5^ plaque forming units (PFU) or a control medium with 5 mg/ml Kanamycin (Wako). From 72 h post-infection, each larva (n = 6) was placed in a 100 mm cell culture dish, round shape of which does not hinder larval locomotory activities and allows the larvae to move continuously. A portion of artificial diet and a piece of black paper were set at the bottom of each dish (Figure 1A). Locomotion was recorded at a 3 s interval using a time-lapse camera (TLC200-pro, Bruno) under light conditions (Figure 1A). We conducted the assay twice as assays #1 and #2.

**Figure 1.**
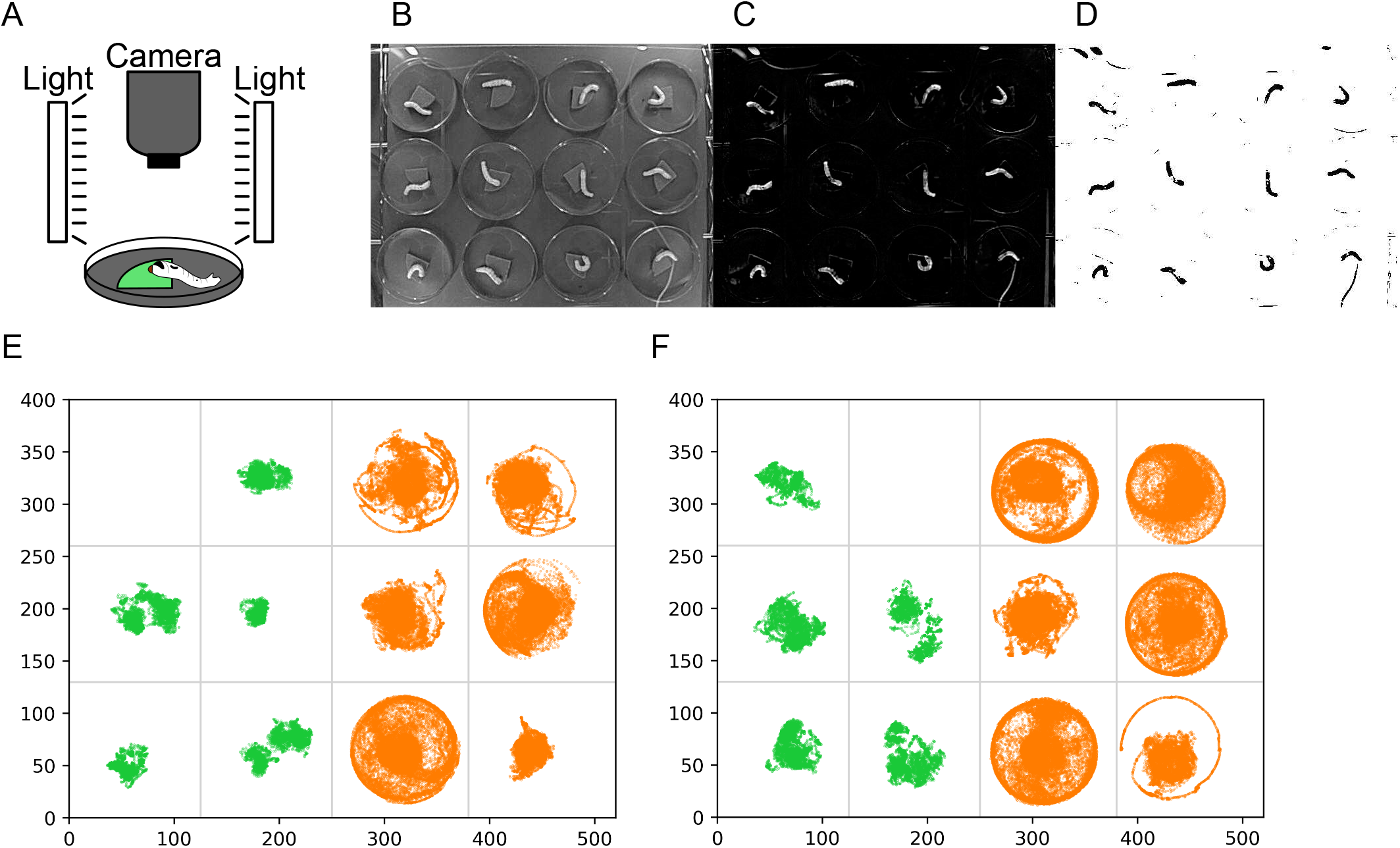
The summary of the behavioural monitoring system. (A) Schematic image of the equipment. The circle and green quadrant represent a cell culture dish and a portion of artificial diet, respectively. (B–D) Image processing. The obtained image was converted to grayscale (B), and the background was removed (C), followed by binarization (D). (E, F) Distribution of larvae for 24 h observation in assays #1 (E) and #2 (F). Each dot indicates the larval position in a frame. Green and orange colours indicate mock- and BmNPV-infected larvae, respectively.

### (c) Video processing

After 24 h of recording, the video was converted to grayscale (Figure 1B). The background image was created as the average intensity. The background was subtracted from all frames in the grayscale video, and then binarization was conducted (Figures 1C and 1D). The processed video was used to determine the larval positions at each frame by calculating the centre of mass of detected particles. These image-processing steps were conducted in Image J software [22].

### (d) Data analysis

The obtained data containing X- and Y-coordinates of the centre of mass were analysed using custom Python scripts. First, we plotted the coordinates onto the original video and checked whether the larval positions were determined appropriately. Based on this analysis, apparently inappropriate points (i.e. dots outside the dish) were identified and removed from the data. Moreover, we found that the positions of some mock-infected larvae were inappropriately mapped because the larvae were so inactive that the method could not create an appropriate background image for these larvae. We excluded the data for these larvae from further analysis. Consequently, we excluded one larva from each assay. The modified data were used for calculating the larval travel distance between two adjacent frames. When multiple points were assigned to a larva in a single frame, we adopted the average of coordinates as the larval position. When the larval position was missing in a frame, the position at its previous frame was used for calculation.

### (e) Quantification of larval locomotion

The total travel distance was calculated as the sum of travel distance between two adjacent frames for 24 h. The accumulated travel distance was also calculated at each time point. Sums of travel distance for each 3 h segment were also calculated. We assumed that a larva exhibited “locomotion” when the average travel distance for 30 s exceeded 0.5 pixels. Moreover, we defined locomotory “continuity” as the number of serial 30 s periods in which a larva exhibited “locomotion.” When the distance of the larval position between two adjacent frames exceeded 0.5 pixels, we assumed the larvae was “moving”; otherwise, the larva was assumed to be “pausing.”

### (f) Statistical analysis

A rank-sum test was used to compare the total travel distance, median locomotory duration, frequency of locomotion, and median travel distance during larval movement between mock- and BmNPV-infected larvae. The sums of 3 h travel distance were compared between mock- and BmNPV-infected larvae using a rank-sum test, followed by Benjamini–Hochberg correction.

## 3. Results

### (a) Overview of larval locomotion over 24 h

Our method successfully traced the locomotion of mock- and BmNPV-infected larvae for 24 h (Supplementary video and Figure 1). Mock-infected larvae generally remained close to a piece of artificial diet showing little locomotion, whereas the locomotory area of BmNPV-infected larvae covered almost the entire area of the dish in both assays #1 and #2 (Figures 1E and 1F). The accumulated travel distance of BmNPV-infected larvae increased continually and more rapidly compared with mock-infected larvae (Figures 2A and 2C). As a consequence, the total travel distance of BmNPV-infected larvae over 24 h was significantly longer than that of mock-infected larvae in two independent assays (Figures 2B and 2D).

**Figure 2.**
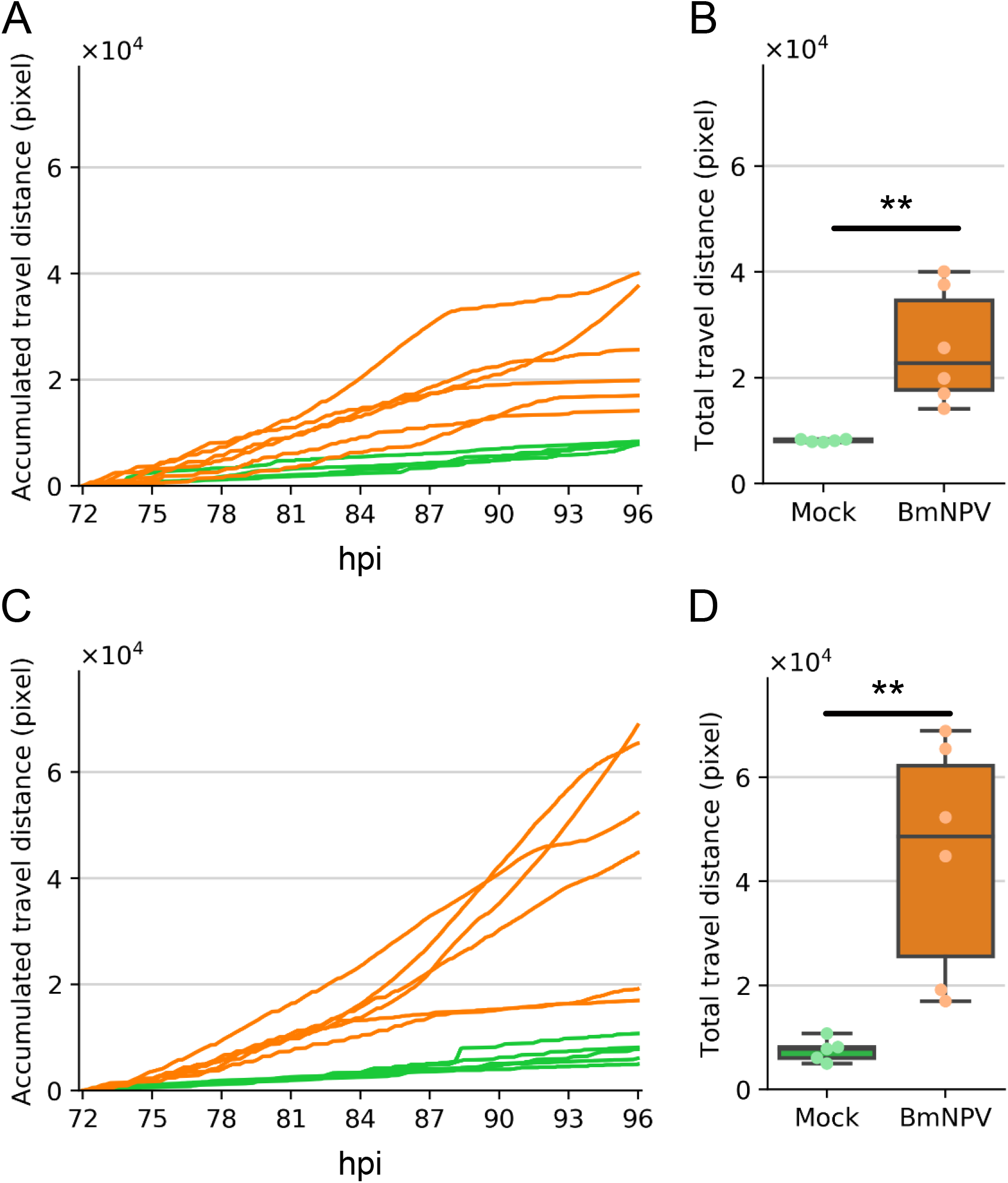
(A) The accumulated travel distance of an individual larva in assay #1. The X- and Y-axes show hours post infection (hpi) and accumulated travel distance at each time point, respectively. The travel distance was measured in pixels. (B) The total travel distance for 24 h in assay #1 shown as a box-and-whisker plot. Boxes show the quartiles, and the whiskers extend to show the rest of the distribution. \*\**p < 0.01* by rank-sum test. (C) and (D) show the accumulated and total travel distance of assay #2, respectively. Green and orange colours indicate mock- and BmNPV-infected larvae, respectively.

### (b) Larval locomotion with 3 s resolution

Larval locomotion of mock- and BmNPV-infected larvae were investigated at a 3 s resolution. We found that mock-infected larvae periodically repeated moving and pausing as described previously [17–19] (Figures 3A, 3C, S1, and S2). This periodic pattern of moving and pausing was also observed in BmNPV-infected larvae, but they were more active than mock-infected larvae (Figures 3B, 3D, S1, and S2).

**Figure 3.**
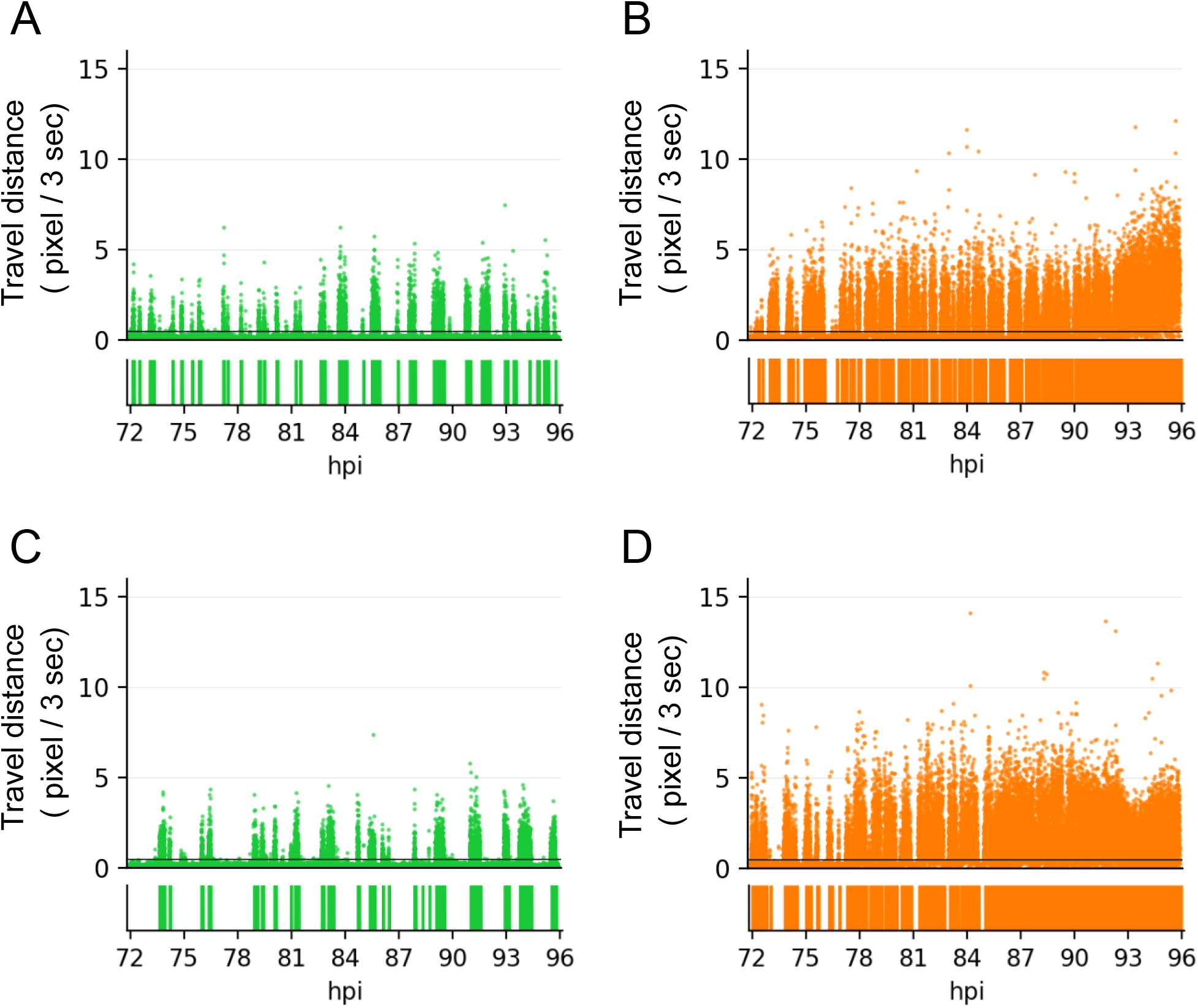
Travel distance between two adjacent frames, which corresponds to 3 s. A representative larva was selected from mock-infected larvae of assays #1 (A) and #2 (C) and the BmNPV-infected larvae of assays #1 (B) and #2 (D). Black lines indicate travel distance equal to 0.5 pixels in 3 s, which correspond to the threshold of moving and pausing. The rectangular diagram below the scatter plots indicates “continuity” of the locomotion, as defined in this study. Green and orange colours indicate mock- and BmNPV-infected larvae, respectively. Data of all individuals are shown in the Supplementary Materials (Figures S1 and S2).

### (b) Quantitative analysis of larval locomotion

The duration of larval locomotion was extended in BmNPV-infected larvae compared with that of mock-infected larvae (Figures S3 and S4). The medians of locomotory duration in BmNPV-infected larvae were ~700 s, which were significantly longer than those of mock-infected larvae (Figures 4A and 4B). In contrast, there was no significant difference between the frequency of continuous locomotion over 24 h of mock- and BmNPV-infected larvae (Figures 4C and 4D). Furthermore, we found that BmNPV-infected larvae moved slightly faster than mock-infected larvae (Figures S5 and S6). The medians of larval locomotory speed during movement were significantly higher in BmNPV-infected larvae than those of mock-infected larvae (Figures 4E and 4F).

**Figure 4.**
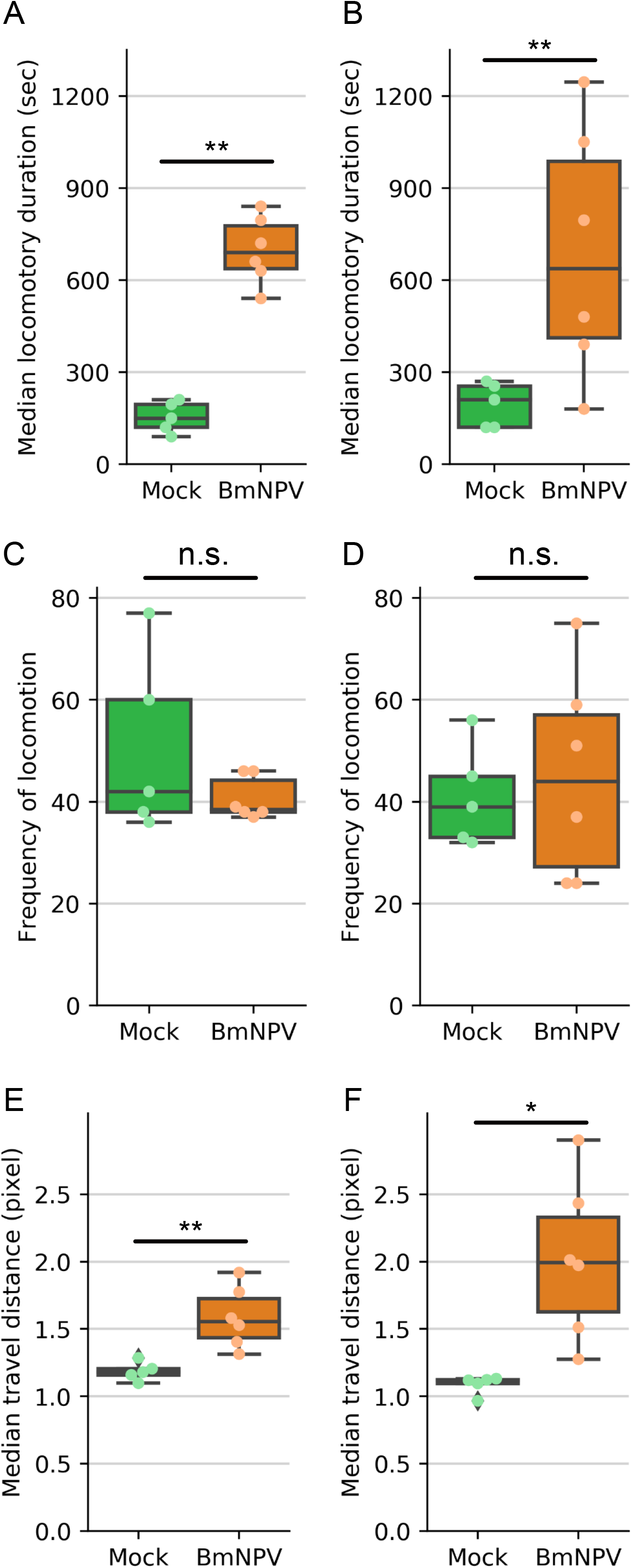
(A, B) Medians of the locomotory duration of assays #1 (A) and #2 (B). (C, D) Frequency of locomotion of assays #1 (C) and #2 (D). (E, F) Medians of travel distance between two adjacent frames during larval movement. Assays #1 (E) and #2 (F). All boxes and whiskers show the quartiles and 1.5 inter quartile range, respectively. Dots indicate values of each larva. \**p* < 0.05 and ** *p* < 0.01 by rank-sum test, respectively.

### (c) Temporal change in larval locomotory patterns

To examine temporal change in larval locomotory pattern, 24 h data were divided into eight 3 h segments and separately analysed. In all 3 h segments, mock-infected larvae stayed in the centre of the dish, where an artificial diet was placed (Figures 5A, 5C, S7, and S8). BmNPV-infected larvae also exhibited food-oriented distribution with slightly larger diffusion during the early time segments (Figures 5B, 5D, S7, and S8). In contrast, BmNPV-infected larvae moved along the edge of the dish and rarely stayed in the centre of the dish in later segments (Figures 5B, 5D, S7, and S8). The sums of travel distance of BmNPV-infected larvae in the segments were generally longer than those of mock-infected larvae, regardless of the infection stages (Figures 5E and 5F).

**Figure 5.**
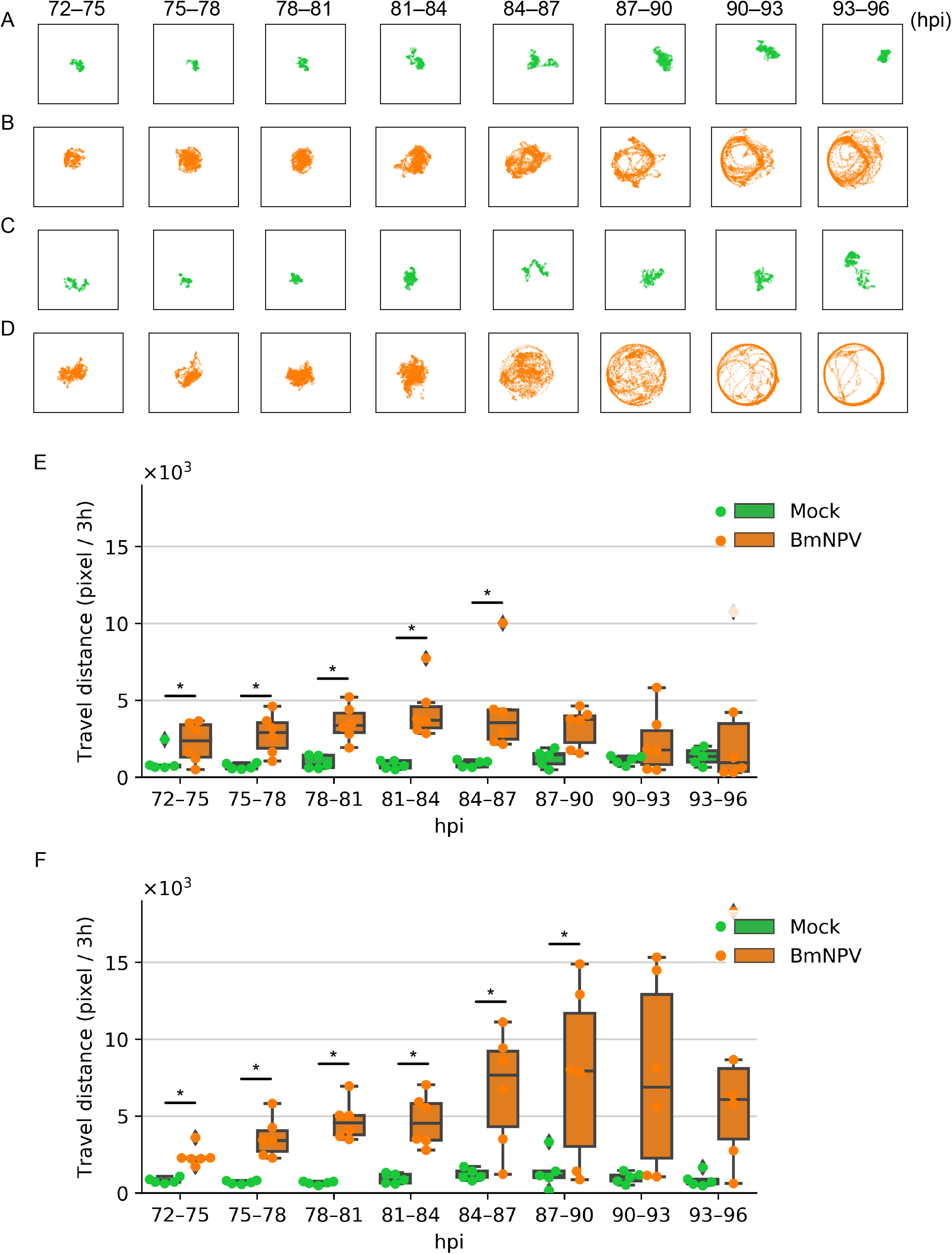
(A–D) Distribution of individual larva for 3 h segments. The time range is shown above the boxes. Each dot indicates the larval position in a frame. A representative larva was selected from mock-infected larvae of assays #1 (A) and #2 (C) and the BmNPV-infected larvae of assays #1 (B) and #2 (D). The individuals are identical to those shown in Figure 3. Data of all individuals are shown in the Supplementary Materials (Figures S7 and S8). (E, F) Sum of travel distance for each 3 h segment for assays #1 (E) and #2 (F). The data are shown as a box-and-whisker plot. Boxes show the quartiles, and the whiskers extend to show the rest of the distribution. Outliers distributed outside of the 1.5 inter-quartile range are indicated as black dots. Coloured dots indicate values of each larva. *Adjustment *p* < 0.05 by the rank-sum test with Benjamini–Hochberg correction. Green and orange colours correspond to mock- and BmNPV-infected larvae, respectively.

## 4. Discussion

We developed a long-term observation system using a time-lapse camera and continuously quantified the locomotory activity of BmNPV-infected larvae over 24 h. Our method enabled us to clarify the difference between the locomotory activity of mock- and BmNPV-infected larvae using a single recording (Figures 1 and 2). Compared with previous methods in which larval locomotion was recorded at multiple time points [11,14,23,24], this method provides a much easier way to quantifying larval locomotory activities during baculovirus infection.

During the continuous observation, mock-infected larvae periodically repeated moving and pausing, which coincides with previous studies (Figures 3A and 3C) [17–19]. BmNPV-infected larvae showed a similar periodic pattern, but the activity level was different from that of mock-infected larvae (Figures 3B and 3D). Quantitative analysis revealed that BmNPV-infected larvae exhibited significantly longer locomotory duration and faster locomotion than mock-infected larvae, whereas the frequency of moving was similar between mock- and BmNPV-infected larvae (Figure 4). These results indicate that BmNPV infection does not induce larval locomotion but increases the locomotory duration and speed, which causes infected larvae to move extremely long distance. Furthermore, we found that both mock- and BmNPV-infected larvae moved close to a piece of artificial diet at the early stage of infection, which indicates that BmNPV-infected larvae maintain a naive locomotory pattern in this infection stage (Figures 5A–D). In contrast, at the late stage of infection, BmNPV-infected larvae moved around the peripheral area of the dish and were rarely found in the centre, (Figure 5B and 5D). This behaviour was not observed in mock-infected larvae (Figures 5A and 5C). These data indicate that BmNPV reprograms the locomotory pattern of its host at a certain time point during the late stage of infection. As larval travel distance was generally longer throughout the late stage of infection, fast and extended locomotion is precedent to the reprogramming of host locomotory pattern and continues independently of this reprogramming (Figures 5E and 5F).

In summary, our study provides a novel insight into BmNPV-induced host manipulation in which BmNPV first enhances the speed and duration of host locomotion, and then reprograms host locomotory pattern, inducing distinct larval behaviour (Figure 6). Currently, baculovirus-induced host manipulation is thought to be biphasic, in which the two phases are divided by whether larval locomotion is horizontal or vertical. These two types of locomotion correspond to ELA (also known as hyperactivity) and CB (also known as tree-top disease), respectively [8,10,12]. Recent studies showed that CB is evoked by phototaxis and that a light signal just before the onset of CB fixes the locomotory direction of the larvae during CB [25,26]. It has also been shown that the larval locomotory direction is irreversible after the critical time point [26], implicating that a reprogramming point of the larval locomotory pattern detected in this study may correspond to the transition point from ELA to CB. As this study did not evaluate vertical movement or phototaxis of *B. mori* larvae, further analysis is required to determine whether the biphasic behaviour observed here is simulates the ELA/CB model. Tentatively, we designate the two phases of the biphasic behaviour “marathon behaviour (MB)” and “lost-child phase (LCP),” which represent the fast and long-lasting locomotion and the reprogramming of the larval locomotory pattern, respectively (Figure 6).

**Figure 6.**
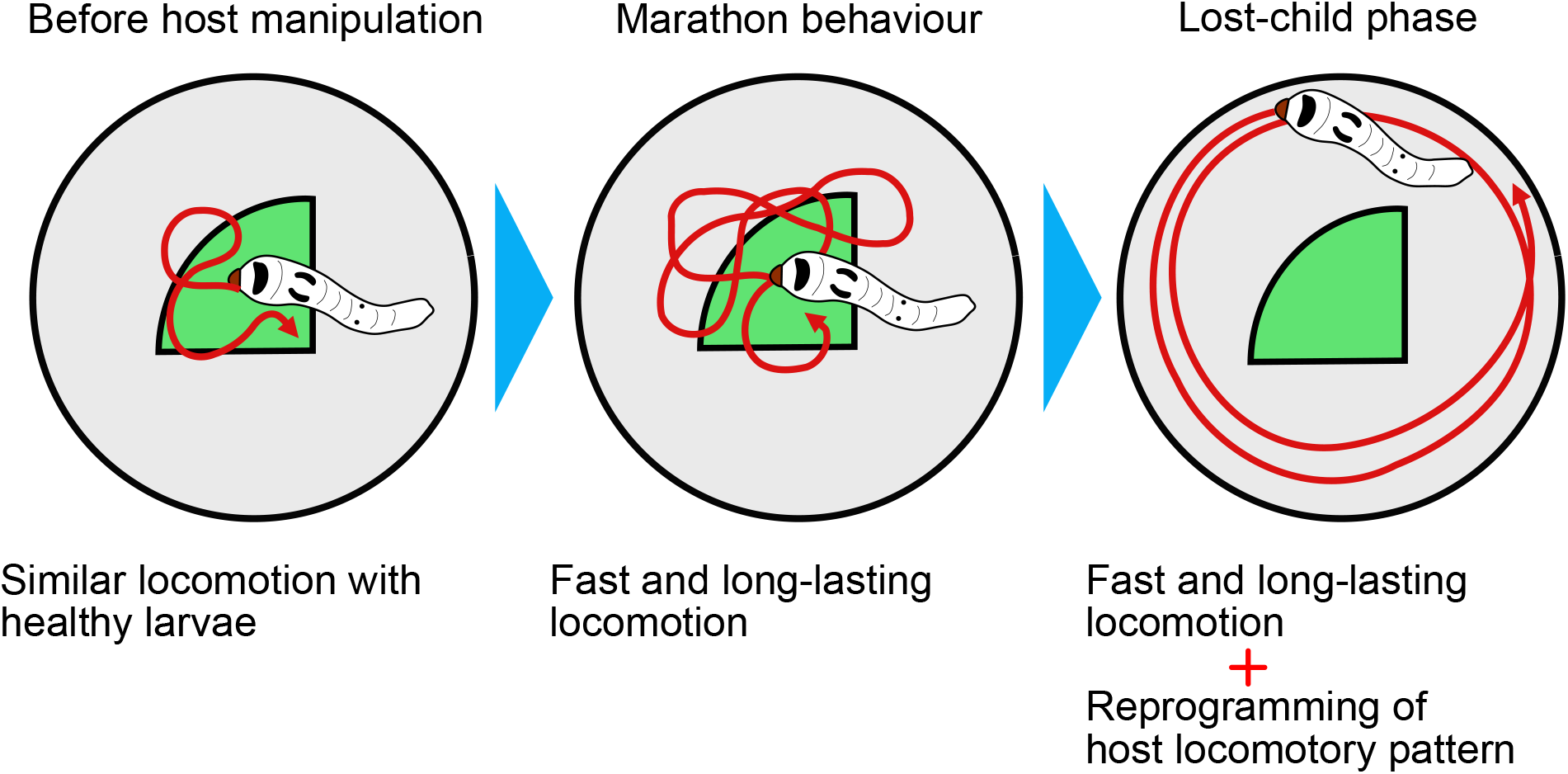
A proposed model for BmNPV-induced host manipulation. The circles and green quadrants represent the cell culture dishes and portions of artificial diet, respectively. Arrows indicate the virtual tracks of larval locomotion.

In this study, BmNPV and *B. mori* larvae were used to assess larval activity during baculovirus infection, but *B. mori* is a highly domesticated insect that exhibits a distinct behavioral pattern compared with its putative ancestor, *B. mandarina* [17]. *B. mandarina* exhibits fast-moving but long-term pausing, whereas *B. mori* repeats moving and pausing over a short period [17]. As BmNPV can infect both *B. mori* and *B. mandarina*, it should be elucidated whether MB and LCP are observed in *B. mandarina* larvae infected with BmNPV, as well as in other host-baculovirus combinations. The continuous observation system developed here is applicable to other insects, although equipment size and observation period should be optimized for the particular insect species. The comparison of host locomotion in various host-baculovirus combinations may clarify the conserved range of MB/LCP model and whether the model is identical to the ELA/CB model, which will highlight underlying mechanisms for the baculovirus-induced host manipulation system.

## Authors’ contributions

H.H. conceived the study and conducted the experiments. H.H. and S.K. analysed the data and wrote drafts of the manuscript.

## Acknowledgement

We thank Takashi Kiuchi and Kenta Tomihara for useful comments. The authors would like to thank Enago (www.enago.jp) for the English language review.

