## Supplemental Figures for "Reprogramming of larval locomotory pattern during baculovirus-induced host manipulation"

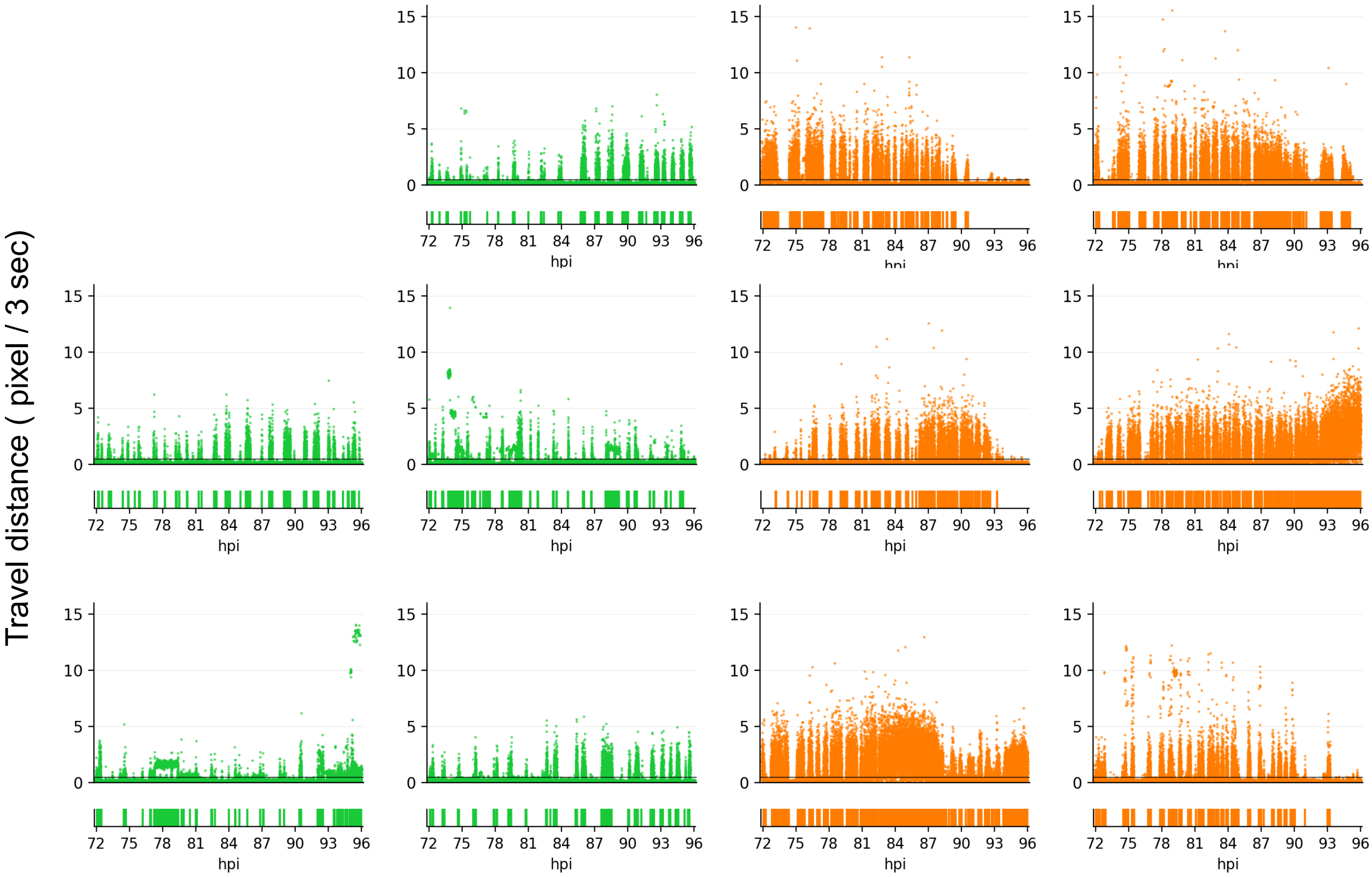


**Figure S1**

Travel distance between two adjacent frames, which corresponds to 3 s for each individual in assay #1. Distribution of individuals corresponds to Figure 1. Black lines indicate travel distance equal to 0.5 pixels in 3 s, which correspond to the threshold of moving and pausing. The rectangular diagram below the scatter plots indicates “continuity” of the locomotion, as defined in this study. Green and orange colours indicate mock- and BmNPV-infected larvae, respectively.

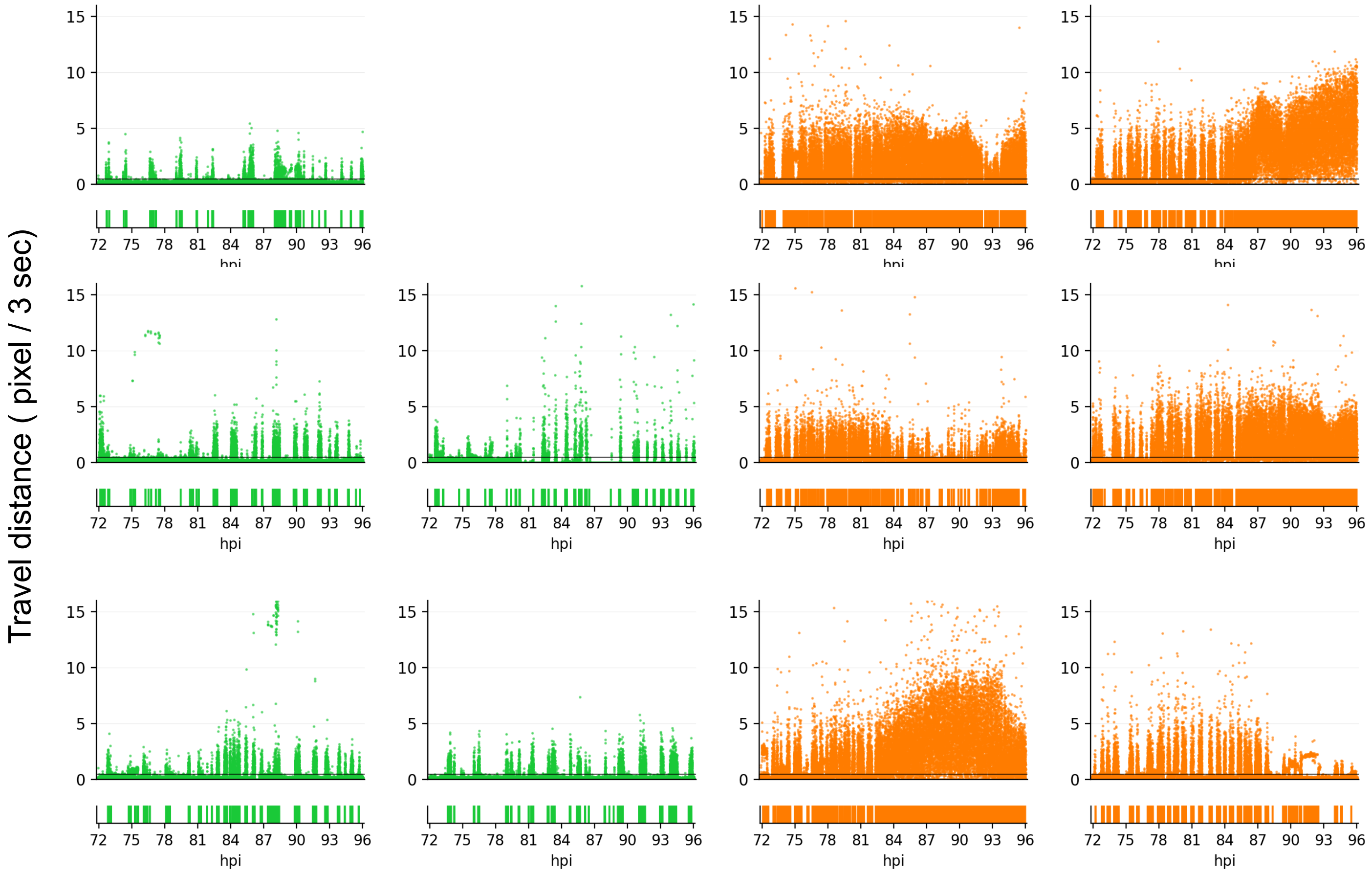


**Figure S2**

Travel distance between adjacent two frames was shown for assay #2 as the same as in Figure S1.

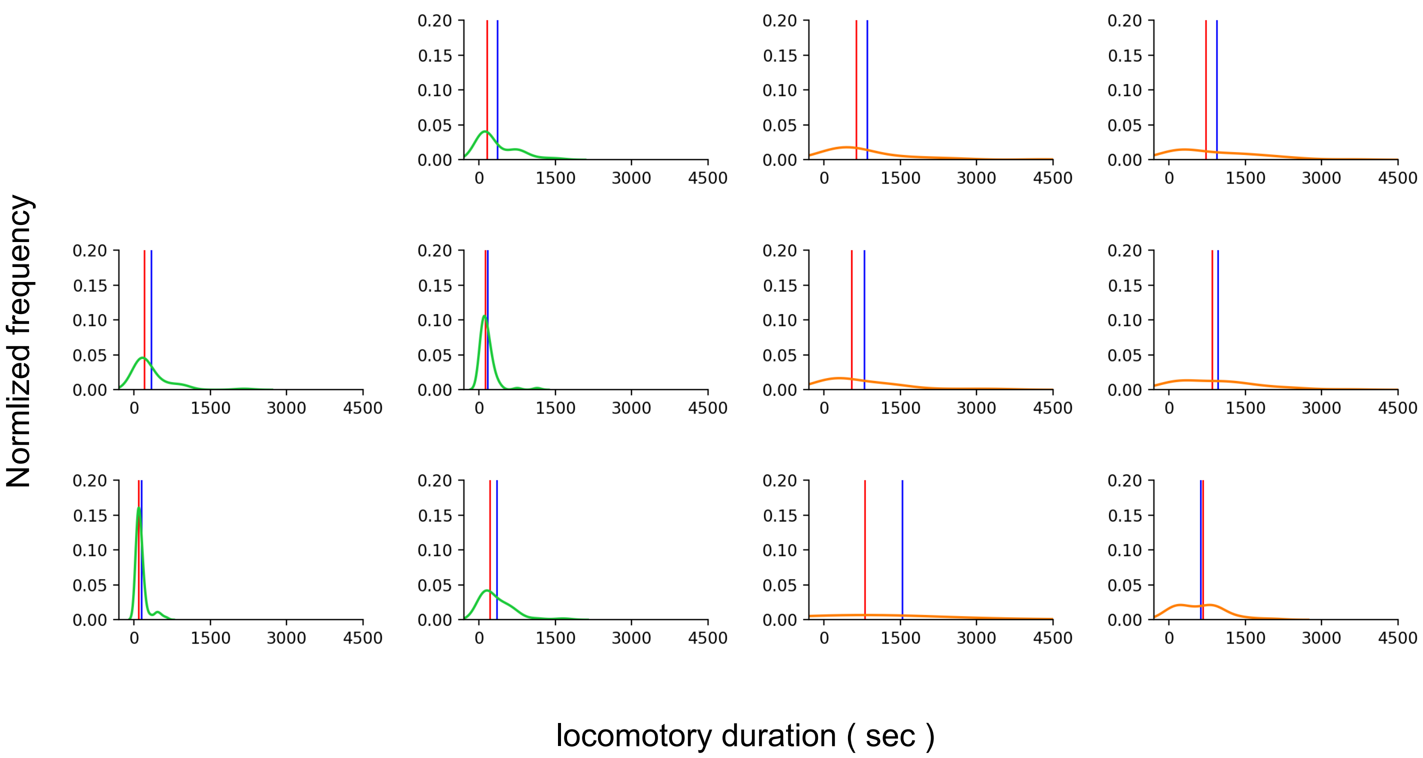


**Figure S3**

The locomotory duration of each larva in assay #1. Each graph indicates an individual larva. Curves represent kernel density estimate. Distribution of individuals corresponds to Figure 1. Green and orange colours indicate mock- and BmNPV-infected larvae, respectively. Red and blue vertical lines indicate the median and mean of locomotory duration, respectively.

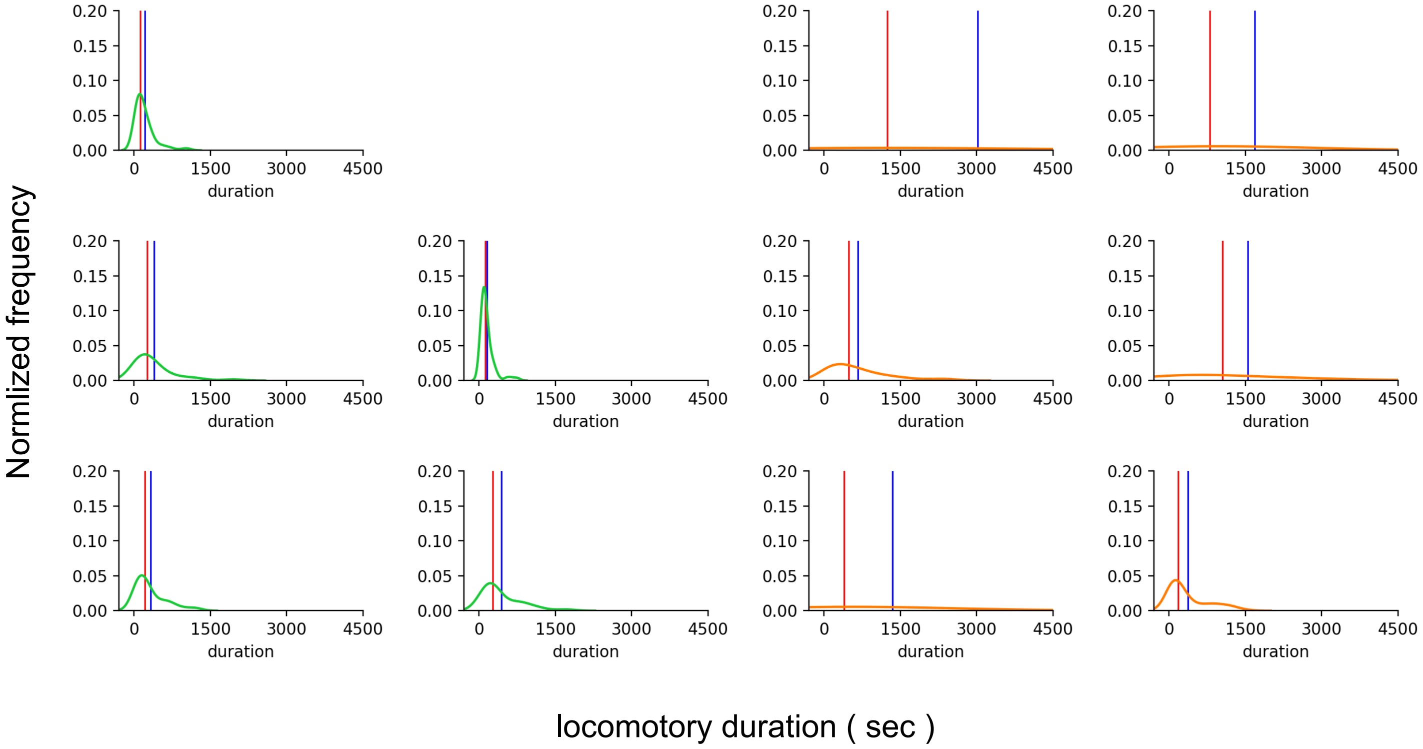


**Figure S4**

The locomotory duration of each larva was shown for assay #2 as the same as in Figure S3.

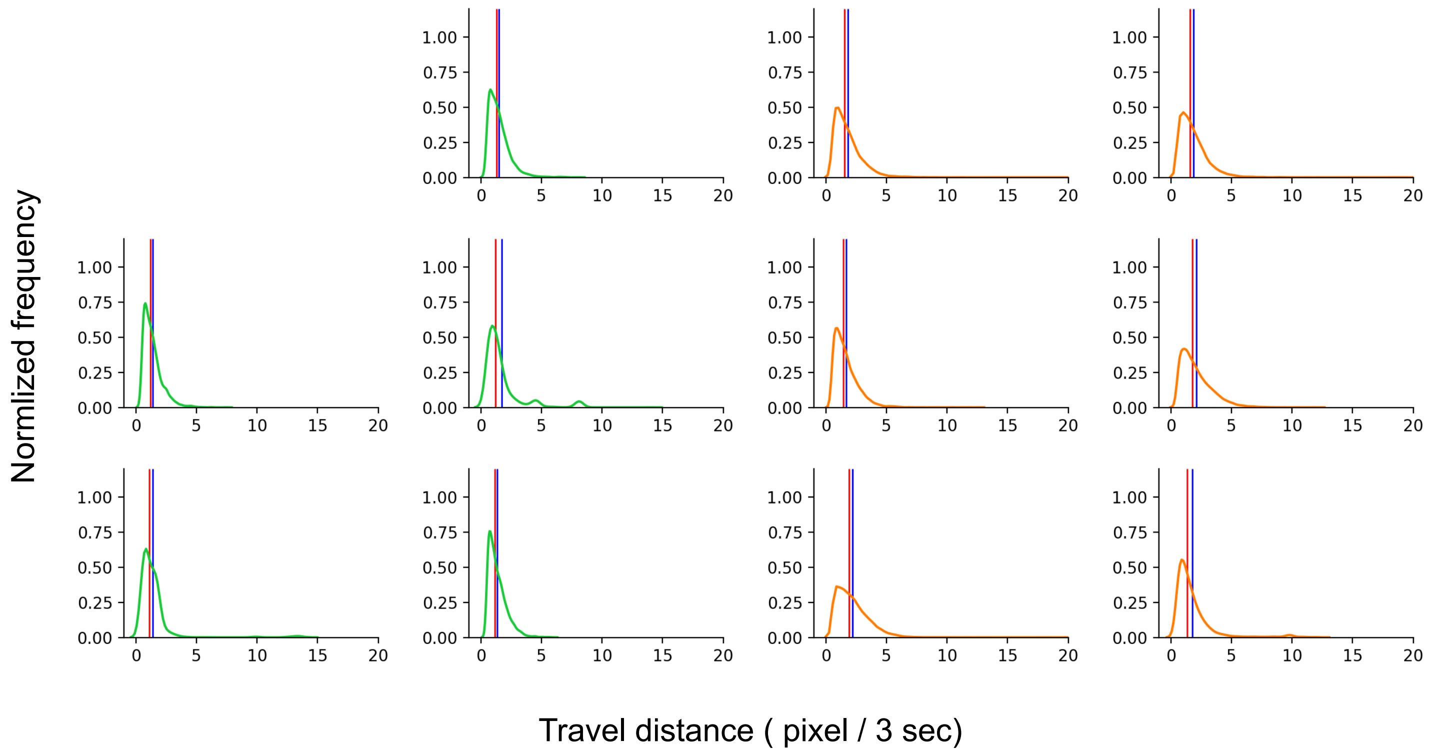


**Figure S5**

Travel distance in each 3 s in assay #1. Each graph indicates an individual larva. Curves represent kernel density estimate. Distribution of individuals corresponds to Figure 1. Green and orange colours indicate mock- and BmNPV-infected larvae, respectively. Red and blue vertical lines indicate the median and mean of travel distance, respectively.

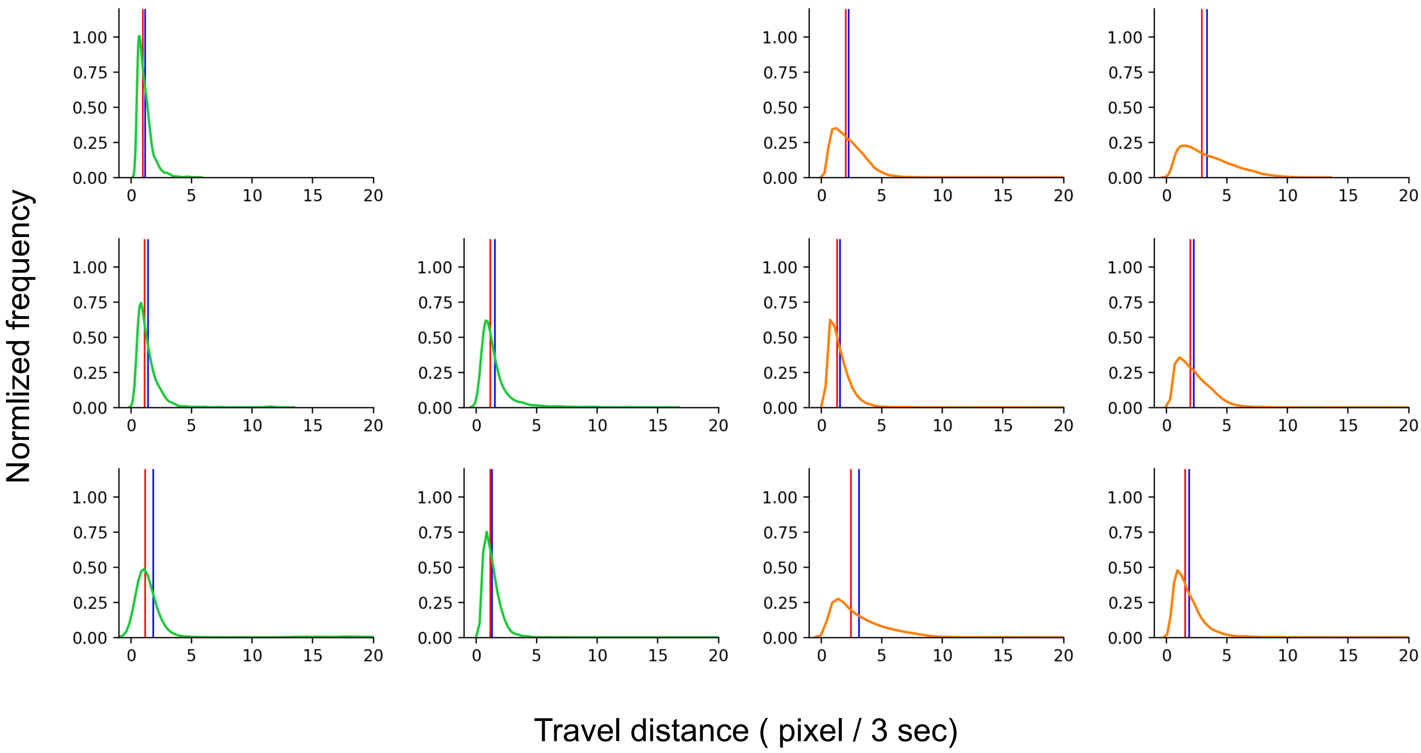


**Figure S6**

Travel distance in each 3 s was shown for assay #2 was shown as the same as in Figure S5.

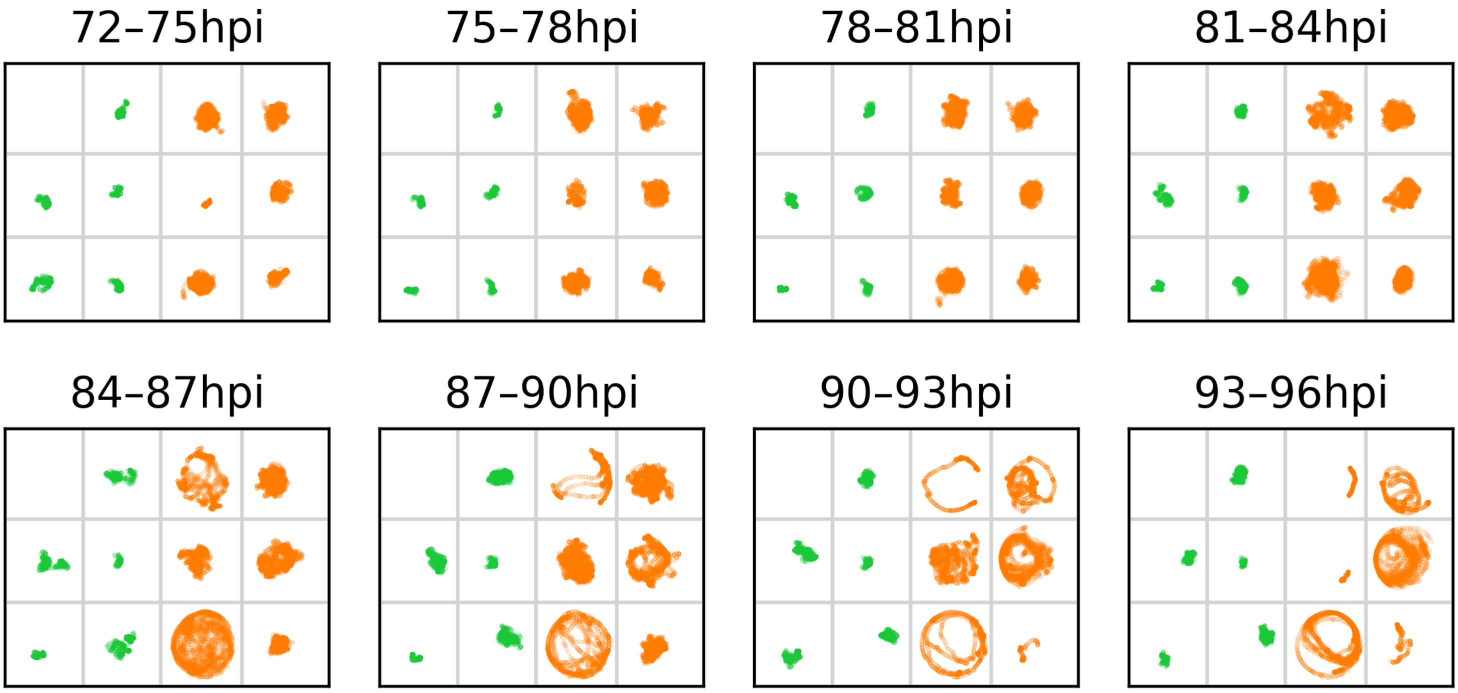


**Figure S7**

Distribution of individual larva for each 3 h segment in assay #1. The time range is shown above the boxes. Each dot indicates the larval position in a frame. Green and orange colours indicate mock- and BmNPV-infected larvae, respectively. Distribution of individuals corresponds to Figure 1.

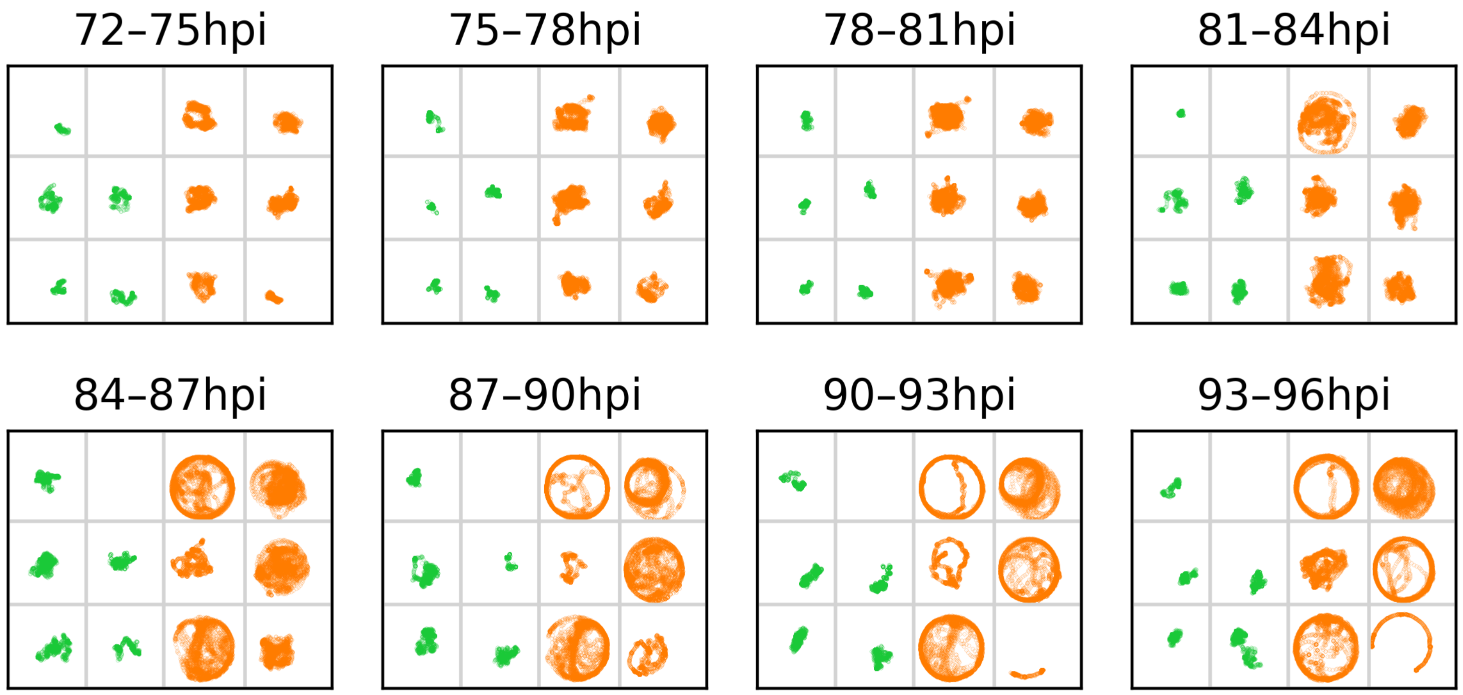


**Figure S8**

Distribution of individual larva for each 3 h segment was shown for assay #2 was shown as the same as in Figure S7.
